## Supporting_information for "Uncoupled evolution of the Polycomb system and deep origin of non-canonical PRC1"

### Supplementary information

### Data and Methods

#### Creation of the proteome database

To profile the occurrence of PRC1 and PRC2 subunits across eukaryotes, we constructed a database comprising the predicted proteomes of 178 eukaryotic species representing the breath of currently-known eukaryotic diversity **(External Dataset S1)**. The representative species were selected on several criteria: i) the overall quality of their predicted proteomes (as measured by BUSCO completeness) (Simão et al. 2015), ii) their phylogenetic position in the eukaryotic tree of life, to ensure an as large as possible diversity within the dataset capturing all currently-defined eukaryotic supergroups and major evolutionary transitions, and iii) the overall interest of the biological community for certain (model) organisms, to ensure relevance and potentially future experimental verification of our inferences.
In order to construct this database, we first combined several previously published eukaryotic proteome selections into a single set. These sets were taken from (Deutekom et al. 2019) the comparative set from (Richter et al. 2022), and the species selection of (Grau-Bové et al. 2022). The resulting dataset comprised 367 species.

As typically splice variants are reported as separate sequences, we filtered splice variants out from the predicted proteomes and translated transcriptomes to produce non-redundant proteomes. Shortly, highly similar sequences from proteomes were clustered using the easy-cluster workflow from the mmseqs2 package (Steinegger and Söding 2017). To evaluate whether the clusters strictly comprised true splice variants, we calculated the percentage identity across the alignment per cluster. First, sequences were aligned using MAFFT (Katoh and Standley 2013). Then, gap-rich regions were excluded from the alignment using the gappyout option of trimal v1.4 (Capella-Gutierrez et al. 2009), as these are possibly the product of in- or excluded exons resulting from alternative splicing. Finally, the percentage identity across all cluster members was calculated from the trimmed alignment with the sident option of trimal v1.4 (Capella-Gutierrez et al. 2009). Clusters containing sequences that all have at least 99% sequence identity in the trimmed alignments to the longest transcript were accepted and the longest transcript was taken as representative. If the percentage identity of a given sequence was below 99% to the cluster representative, the clustering was deemed spurious and the outlier sequences were considered as non-redundant sequences.

BUSCO (v5.2.2; eukarya_odb10) was used to determine the quality of the proteomes of these species. Furthermore, single-copy BUSCO orthologs found in at least 75% of all species were selected and taken as marker genes for the construction of a phylogenetic tree, yielding a comprehensive resolution of the diversity in this combined set. The phylogenetic tree was inferred using IQTree (v2.2.0) using the LG+F+R15 substitution model . The resulting phylogeny was visualised in iTOL, and BUSCO scores were plotted on these allowing for manual selection of species at key phylogenetic positions with the highest available quality proteomes across the relevant taxa (Letunic and Bork 2021). With a total of 178 species, the final set is computationally tractable while also allowing for maximum diversity and relevance to understand the evolution of proteins across eukaryotic lineages, such as the subunits of PRC1 and PRC2.

#### Curation of HMM profiles

Similar to recently published studies, our phylogenetic trees were initially annotated with Pfam domains (v.35) (Mistry et al. 2021) to delineate orthologous groups (Grau-Bové et al. 2022; Sharaf et al. 2022). Sequences however, reported no domains with default Pfam (v.35) HMM models, while these sequences clustered within our orthologous groups (Mistry et al. 2021). Alignments as well as Alphafold 2 predicted (Varadi et al. 2022; Mirdita et al. 2022; Jumper et al. 2021) structures clearly showed the presence of e.g. a RAWUL domain. Custom HMM models were therefore created with the HMMbuild tool of the HMM package (version HMMER 3.3), based on alignments, resolved 3D structures, and Alphafold 2 predicted structures (Varadi et al. 2022; Mirdita et al. 2022; Jumper et al. 2021) **(External Dataset S2)**. This method, together with the highly diverse set of eukaryotic proteomes allowed us to delineate a more detailed view of PRC1 and PRC2 evolution compared to other studies. This argued that previous studies utilizing these available/deposited profiles to detect homologs and delineate orthologs were therefore likely unable to detect similar amounts of domains as in this study.

#### Protein specific analysis strategies

##### *EZH and RBBP*

For EZH and RBBP, analyses were performed as described in the 'materials and methods' section of the main text. For RBBP, we did not annotate the sequences in the phylogenetic trees with domain annotations because the other criteria sufficed to determine the orthologous group.

##### *RING1 and PCGF*

In order to identify orthologs of RING1 and PCGF, initial analyses demonstrated the need for an iterative approach. First, phylogenetic trees were annotated with independent function predictions (eggnog) and predicted domains based on HMM searches with the 'hmmsearch' tool from the HMMER package (http://hmmer.org/, HMMER 3.3), with profiles from Pfam (v.35) (Huerta-Cepas et al. 2019; Mistry et al. 2021). Sequences were considered to be orthologs if they clustered in the orthologous group, and both function (eggnog) and domain (Pfam) predictions were complete, or if both domains in the 3D structures that we predicted with AF2 were detected (Varadi et al. 2022;

Mirdita et al. 2022). From this set of orthologs, a new MSA was created with MAFFT v.7.490 (settings genafpair, maxiterate 1000) (Katoh and Standley 2013). Alignments were then checked for the presence of RING and RAWUL domains **(External Dataset S7)**. HMM profiles were created of these domains with the 'hmmbuild' tool from the HMMER package (http://hmmer.org/, HMMER 3.3) **(External Dataset S2)**. This was done separately for both RING and PCGF. With these new profiles, more instances of domain conservation were detected in our sequences. Sequences were deemed orthologs if both domains were present. Otherwise, they were deemed putative orthologs.

***EED***

Similar to RING1 and PCGF, our phylogenetic inference of EED demonstrated the need for another approach. Pfam (v.35) domain models were again not sufficient in detecting all WD40 domains, and eggnog annotations revealed that a monophyletic cluster that initially was considered to the orthologous group according to our criteria, also contained other proteins. In addition, the initial homology search did not result in the detection of the described EED orthologs in the Ciliates *Paramecium tetraulia* and *Tetrahymena thermophila* (Wang et al. 2022; Xu et al. 2021). Therefore, HMM models were again created with the 'hmmbuild' tool from the HMMER package (http://hmmer.org/, HMMER 3.3) based on the full length sequences of these Ciliates, and then a local HMM search was performed **(External Dataset S2)**. From this output, potential EED hits of Ciliates were selected and added to the alignment. Then the initial phylogeny pipeline as described in 'materials and methods' of the main text was re-run. Additionally, an iterative approach was performed where orthologs were first selected manually. These orthologs were accordingly aligned with MAFFT v.7.490 (settings genafpair, maxiterate 1000) (Katoh and Standley 2013) and from this MSA, HMM profiles were built with the 'hmmbuild' tool from the HMMER package (http://hmmer.org/, HMMER 3.3) based on coordinates in (Chammas et al. 2020), and used to annotate our trees **(External Dataset S7)**. Even now, these profiles did not detect all 7 WD40 domains in the sequences of our cluster **(Fig.S1E)**. Yet, it allowed us to define a more precise orthologous group. A clade with a support value of 87 was eventually considered to be the orthologous group **(Fig.S1E)**. This decision was based on the quality of eggnog annotations, and domain annotations, and on the knowledge of the diverged EED sequences in Ciliates which are orthologs according to the experimental data (Wang et al. 2022; Xu et al. 2021; Huerta-Cepas et al. 2019).

***SUZ12***

For the highly divergent subunit SUZ12, a phmmer search was performed with the HMMER package (http://hmmer.org/, HMMER 3.3 and full length significant hits were selected as orthologs **(External Dataset S4)**. Accordingly, the alignments were manually checked for the presence of representative domains. Then, a full length HMM profile was created based on the selected sequences and another HMM search against our local proteome database was performed. Significant hits (bitscore = 20) were inspected, and aligned with MAFFT v.7.490 (settings genafpair, maxiterate 1000) (Katoh and Standley 2013) and from this MSA, HMM profiles were built with the 'hmmbuild' tool from the HMMER package (http://hmmer.org/, HMMER 3.3) based on coordinates in (Chammas et al. 2020) . No other proteins with a VEFS-box besides SUZ12 were observed in the homology search. Similarly, no other proteins with a VEFS-box were found in other literature. Therefore, the presence of SUZ12 was inferred based on the presence of a VEFS-box. All sequences were manually checked for VEFS-box presence. If a sequence contained the VEFS-box we determined it to be an ortholog. In any other case, sequences were not included in our orthologous group.

***CBX and RYBP***

For CBX and RYBP, orthologs were manually selected and full length HMM profiles of these sequences were created with the 'hmmbuild' tool from the HMMER package (http://hmmer.org/, HMMER 3.3). Accordingly, HMM searches against our local database with these profiles was performed **(External Dataset S3)**.

***SAM-domain containing accessory subunits***

For SAM-domain containing accessory proteins, an online psiblast on MPI toolkit (https://toolkit.tuebingen.mpg.de/) (Gabler et al. 2020; Zimmermann et al. 2018) was performed with the SAM domain of *Homo sapiens* PHC **(External Dataset S4)**. A HMM profile was then created of the MSA output with the 'hmmbuild' tool from the HMMER package (http://hmmer.org/, HMMER 3.3). Subsequently, local HMM searches against our local proteome database were performed with this profile and the Pfam (v.35) profile of MBT. Relevant hits (SAM (bitscore = 50), MBT (bitscore = 25)) were retrieved, and domains were aligned using MAFFT v.7.490 (settings localpair, maxiterate 1000), and processed as mentioned in 'materials and methods' of main text **(External Dataset S4**. After the detection of the L3MBTL-associated orthologs in the unicellular relatives of fungi and animals, the SAM domains from these homologs to our trees were manually added to our trees, as initially they were not included in our phylogeny due to their lower bitscore values than our threshold.

### External datasets

**Please find the datasets described below on figshare:**

https://figshare.com/articles/dataset/Supporting_Information_associated_with_bioRxiv_manuscript_Uncoupled_evolution_of_the_Polycomb_system_and_deep_origin_of_non-canonical_PRC1_/22548865

**Dataset S1**

Proteome database. Table containing all eukaryotic species information of which sequences are used to construct the phylogenetic trees in SI Fig. S1A-H.

**Dataset S2**

Manually curated HMM of RING1, PCGF, EZH, RBBP, EED, and SUZ12.

**Dataset S3**

Materials that were used for the structural alignment. Includes TM align files, HHpred outputs and pdb files of excised C-terminal ß-hairpins.

**Dataset S4**

Materials and Data that were used for the homology analyses of CBX, PHC, RYBP, and SUZ12. For CBX and RYBP, HMM profiles and HMMsearch outputs are provided. For PHC, SAM and MBT HMM profiles and searches, and newick files of our phylogenetic trees are provided. For SUZ12, HMM profile and multiple sequence alignment are provided.

**Dataset S5**

Alphafold structures of putative sequences of RING and PCGF and the newly identified RYBP orthologs.

**Dataset S6**

Blastoutputs of RING, PCGF, EZH, EED, and RBBP.

**Dataset S7**

Multiple sequence alignments used to build phylogenetic trees and newick files for EZH, EED, RBBP, RING and PCGF.

**Dataset S8**

Fasta sequence files of our orthologous groups of the core subunits of PRC1 and PRC2, and newly identified ncPRC1 accessory subunits.

### Figures

#### Figure S1 (Below). Phylogenetic trees of core proteins PRC1 and PRC2

To manually delineate orthologous groups from the phylogenetic trees, trees were annotated with independent function predictions (eggnog) and predicted domains based on HMM searches with our self-curated HMM profiles (Huerta-Cepas et al. 2019). Details on inference methods can be found in **SI Data and methods.** Sequences were obtained from our local proteome database. The first 6 letters of the protein name indicate the species as listed in **External dataset S1**. Clades that we determined to be the orthologous group are represented in red. All trees are plotted as circular trees with bootstrap values and eggnog and domain annotations. For F and G, we plotted the unrooted tree and highlighted the branches of the newly found L3MBTL-associated orthologs in green.

(A) RING

(B) PCGF

(C) EZH

(D) EED

(E) RBBP

(F) SAM

(G) MBT

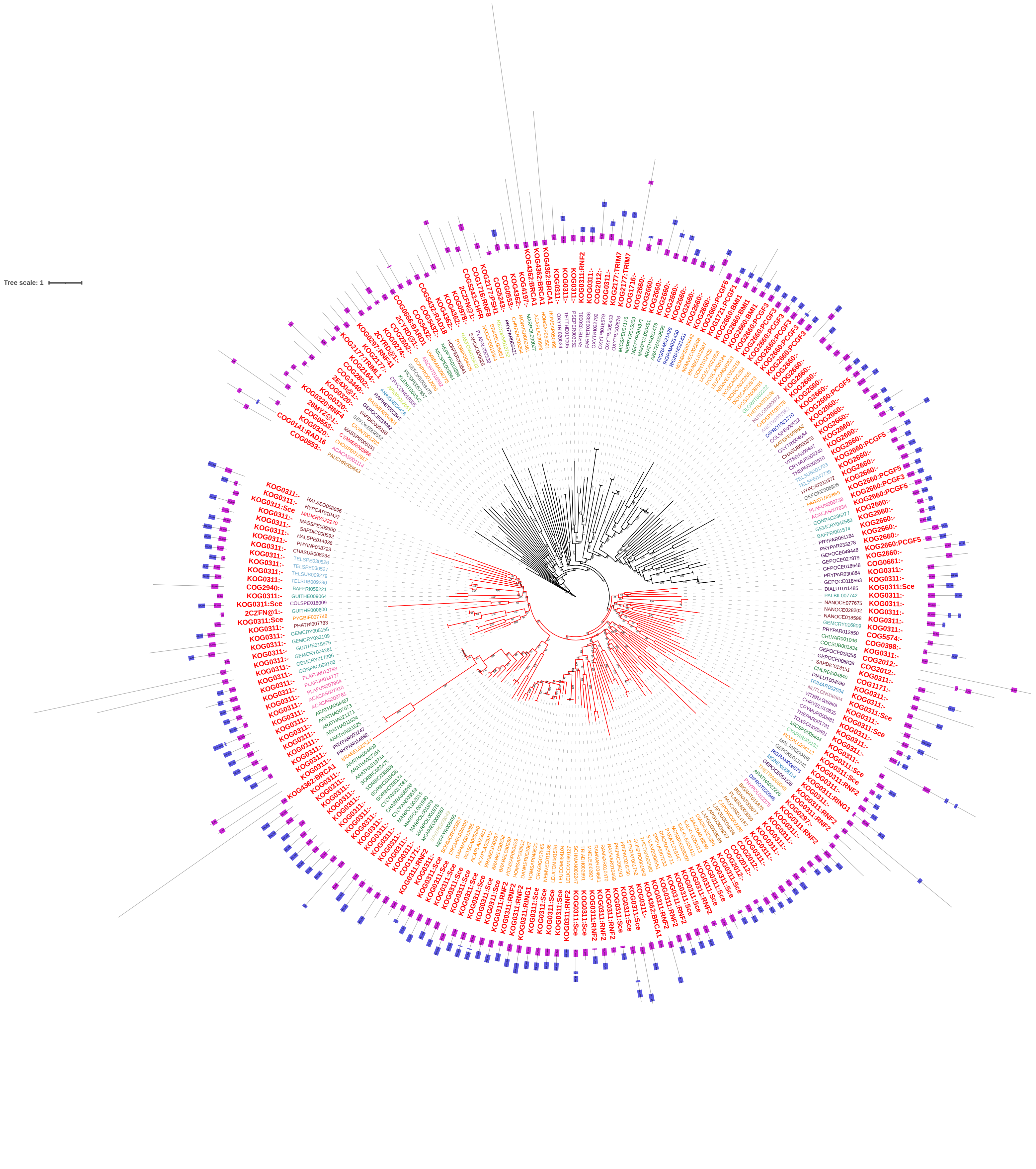

Fig.S1A. Phylogenetic tree of RING1

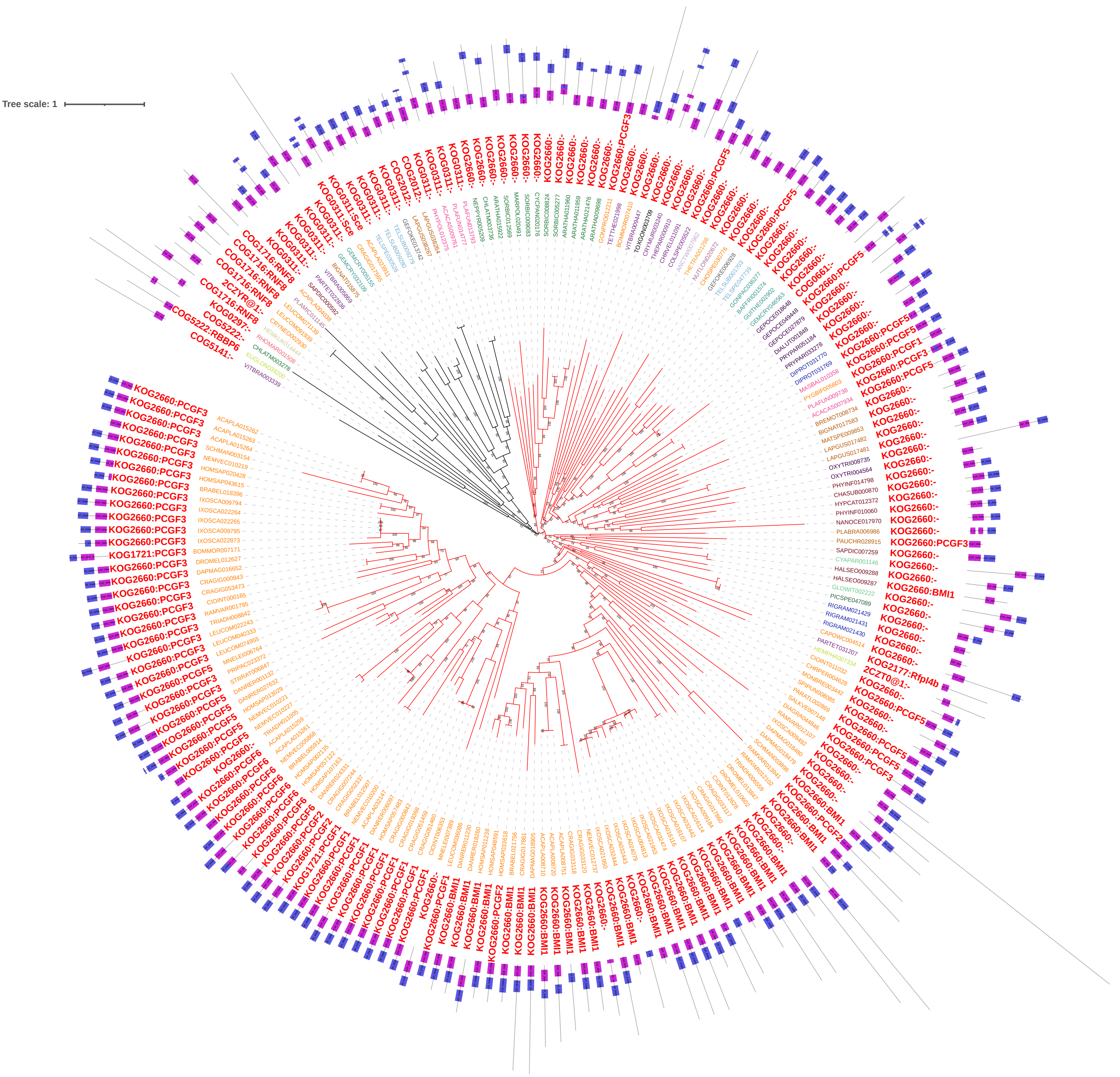


Fig.S1B. Phylogenetic tree of PCGF

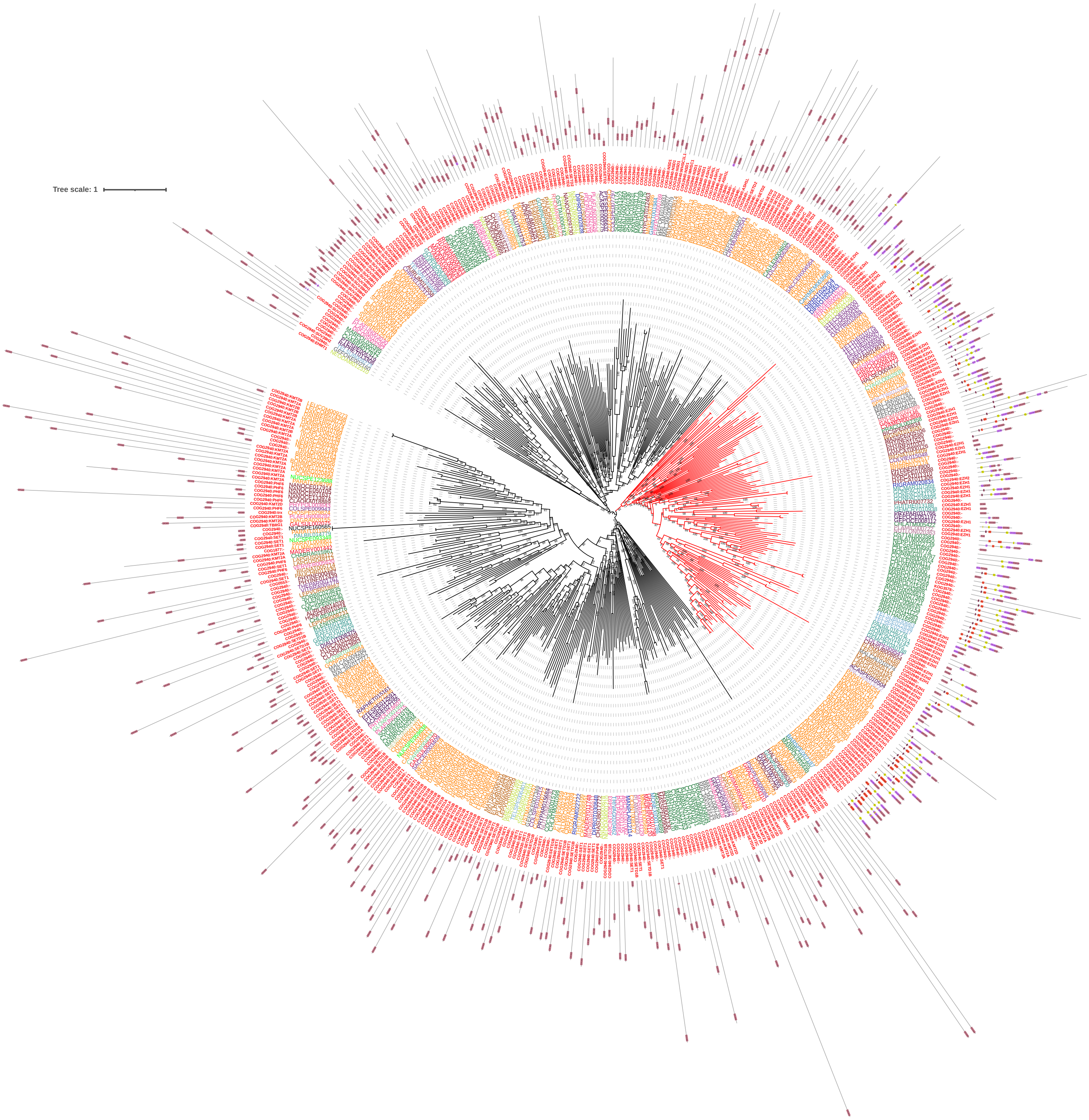


Fig.S1C. Phylogenetic tree of EZH

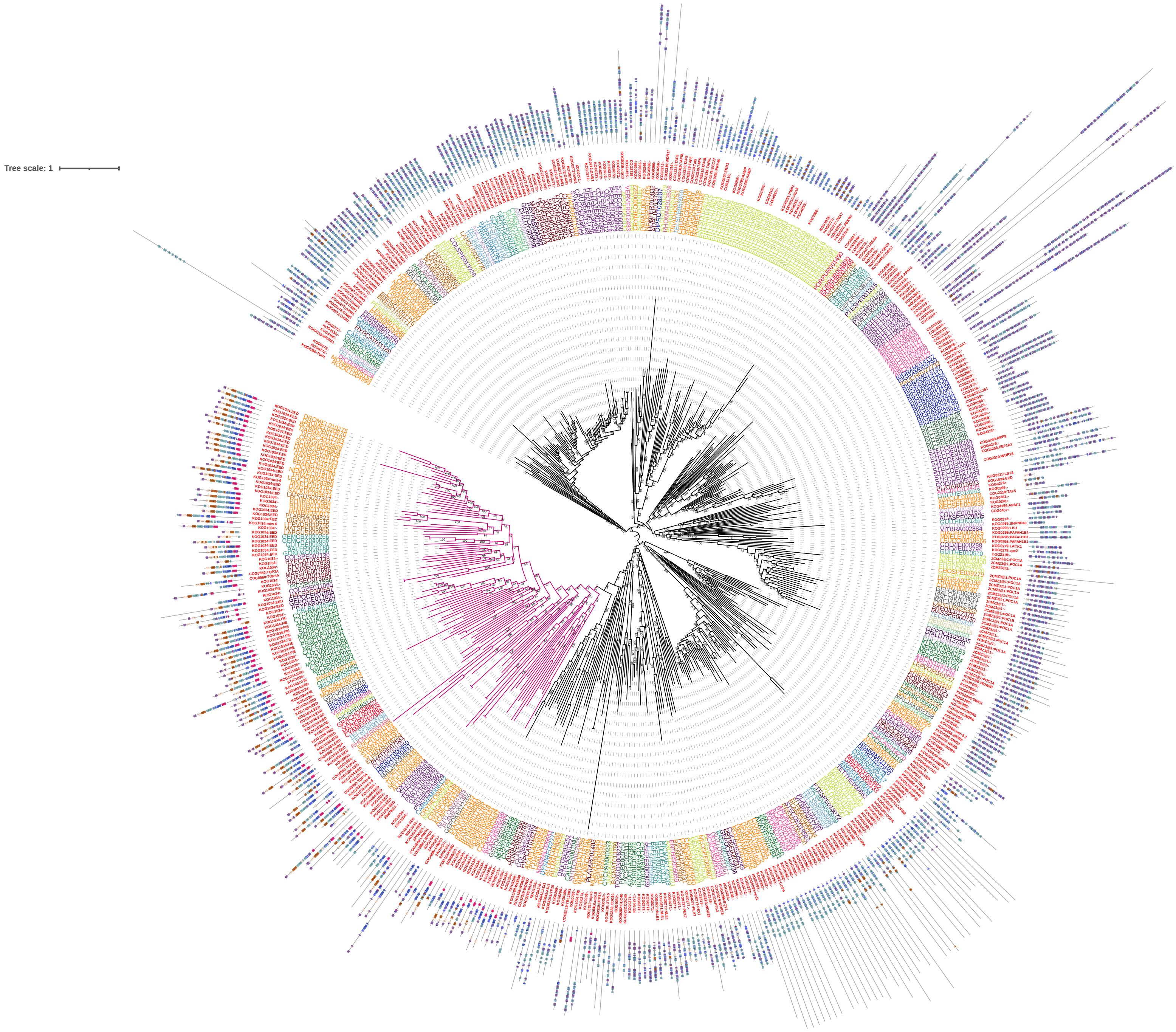


Fig.S1D. Phylogenetic tree of EED

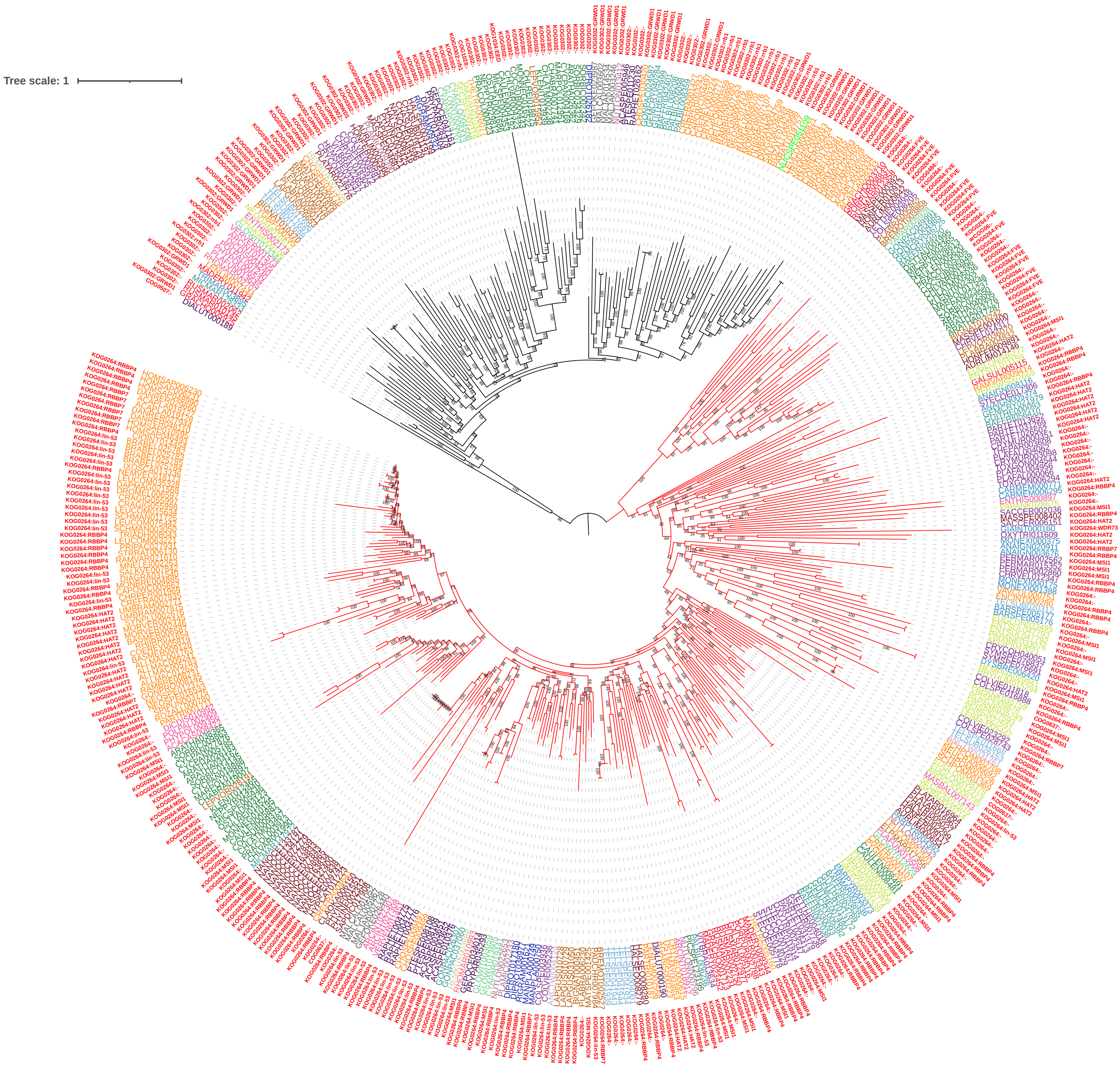


Fig.S1E. Phylogenetic tree of RBBP

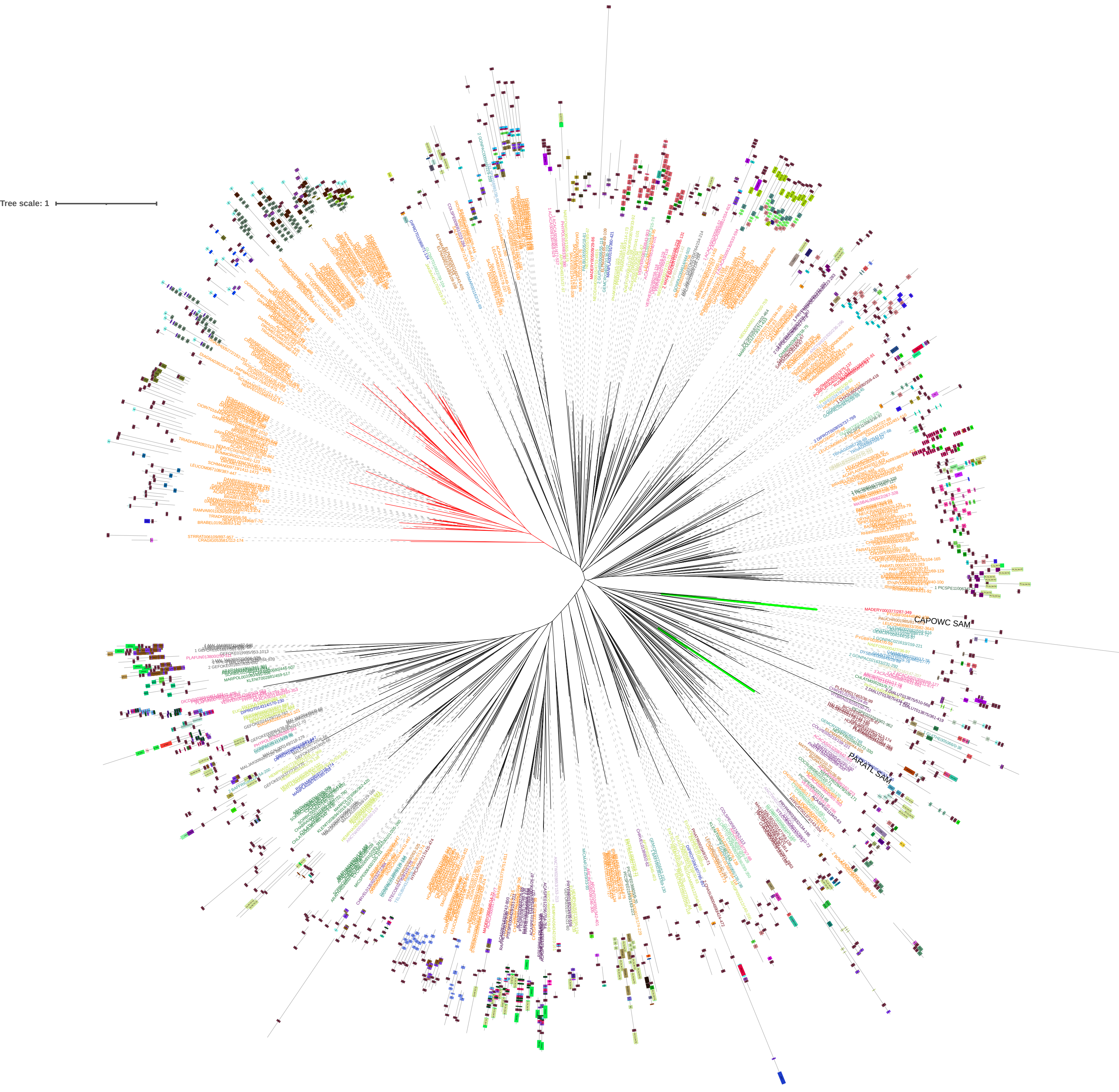


Fig.S1F. Phylogenetic tree of SAM

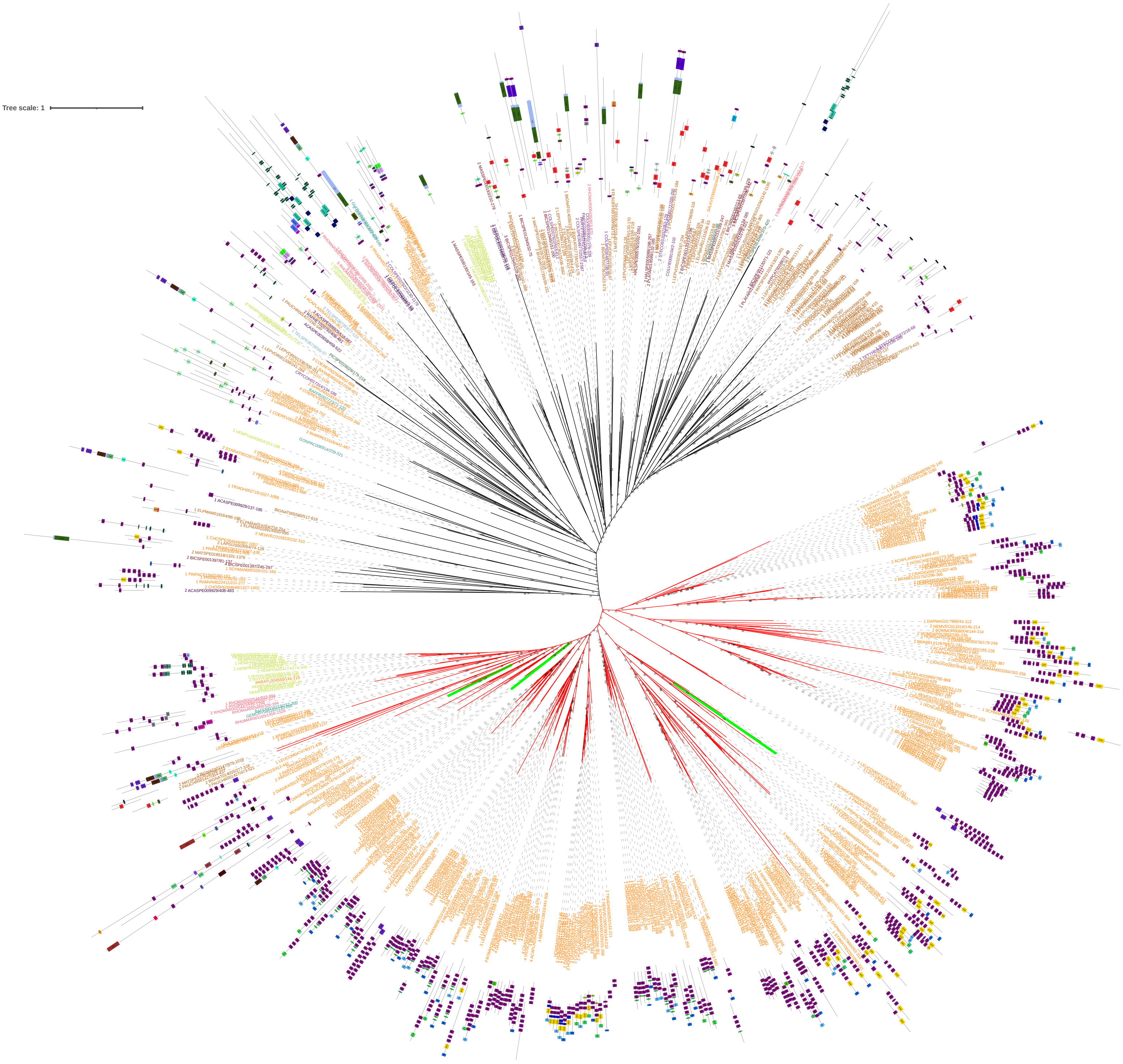


Fig.S1G. Phylogenetic tree of MBT

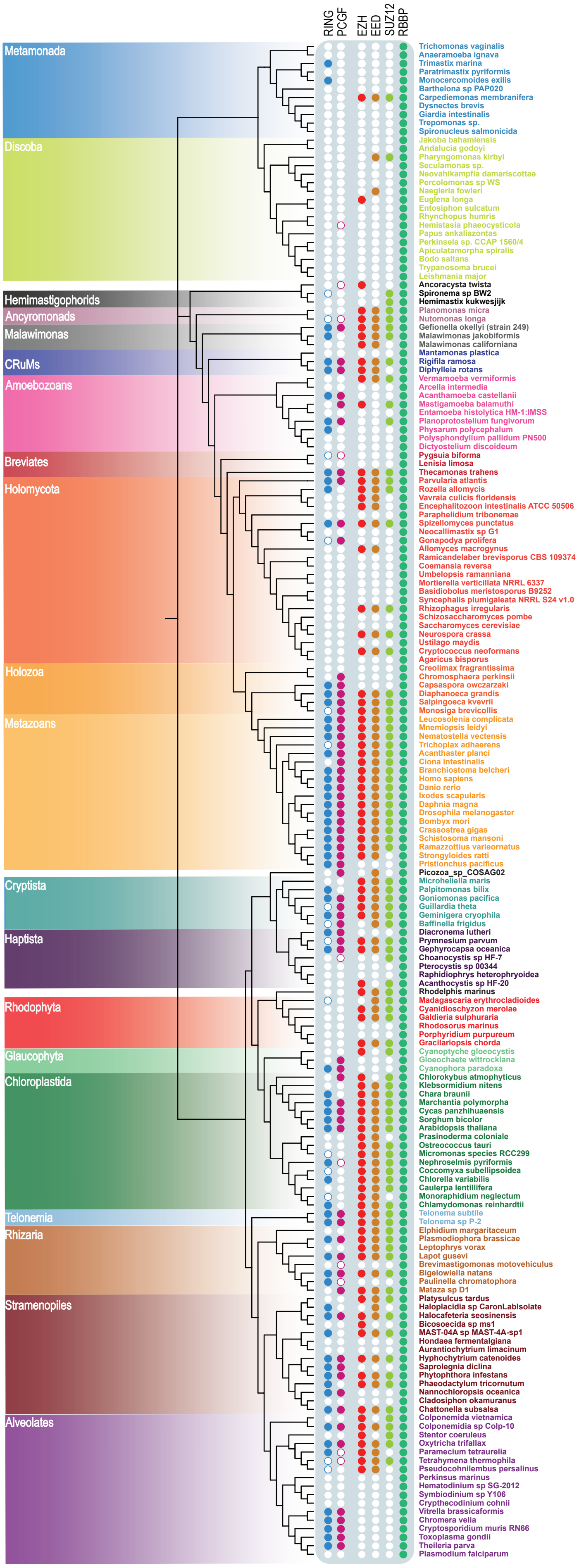


Fig.S2. Phylogenetic profiles of core proteins PRC1 & PRC2. Shown are the phylogenetic profiles of PRC1 & PRC2 core subunits of all independent species included in our analysis

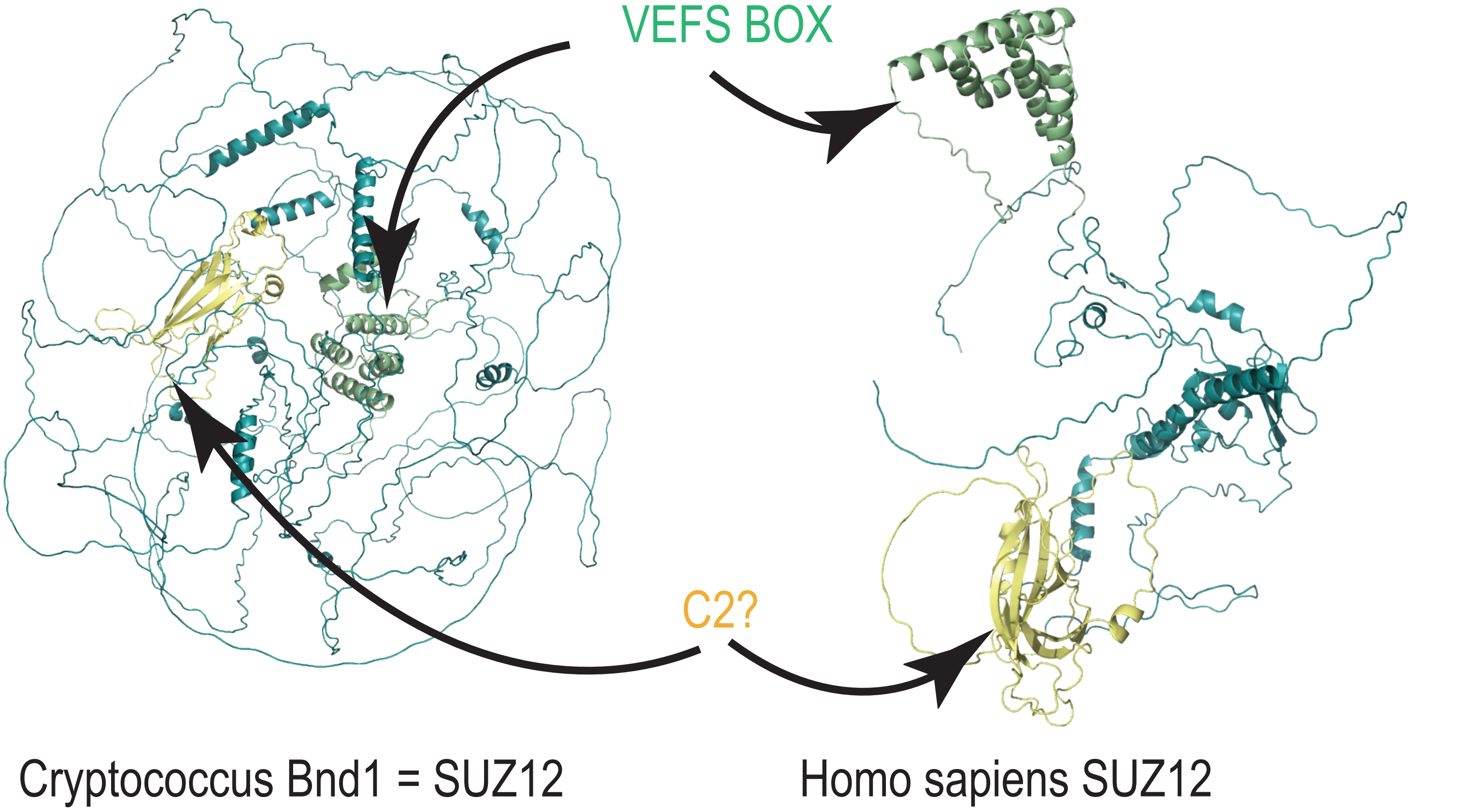


#### HHPRED VEFS BOX

##### Alignments

Template alignment | Template 3D Structure | PDBe

1. **5HYN_S Polycomb protein SUZ12; chromatin modification complex, transferase; HET: ZN, SAH, M3L; 2.95A {Homo sapiens}**
Probability: 98.39%, E-value: 9.4e-7, Score: 64.51, Aligned cols: 82, Identities: 13%, Similarity: 0.126, Template Neff: 8.5

```
Q ss_pred               EeecCccceeehHHhhhhhhcCCCCccCCCcchHHHHHHhhhhhhcCCcHhhHHHhhheeecCCCCHHHHHHHHHHH
Q Q_8062480        33   QHHLSHKRRFLACCWNRWVHQKGPIPAYNKAGYLKHWLDSYGIIMCRAKLSYEVKDYLHLHFQNRHIPLTDYVGACQYWN   112 (126)
Q Consensus        33   ~~Hls~krRFlaCcWNRWV~qkGPIp~~n~~~y~~~fi~~Yg~iM~rA~L~~EVrD~L~i~~~~k~I~l~~~~~a~~~wn   112 (126)
                        ...++++++-++.+||+++...++++...-+.-+..|+++++..+.+.+|-.+..-+|..+++...|+-.++..++++.+
T Consensus        47   ~~dv~~~Ek~fm~lWN~fi~~~~~~~~d~~~~~~~~~~F~~~~~~~~l~~~~l~~~f~~hl~~L~~~~~l~~~~~~~~~l~~~~   126 (129)
T 5HYN_S           47   FSDVNEGEKEVMKLWNLHVMKHGFIADNQMNHACMLFVENYGQKIIKKNLCRNFMLHLVSMHDFNLISIMSIDKAVTKLR   126 (129)
T ss_dssp               CSSSCHHHHHHHHHHHHHHHHHTCCSSTHHHHHHHHHHHHHHHHHHHHTTCHHHHHHHHHHHHHHTTCSCHHHHHHHHHHH
T ss_pred               cCCCCHHHHHHHHHHHHHHHHHcccCchhhHHHHHHHHHHHHHHHHHHhhHHHHHHHHHHHHHHHHCCCCCHHHHHHHHHHH

Q ss_pred               HH
Q Q_8062480       113   HL    114 (126)
Q Consensus       113   ~i    114 (126)
                        ++
T Consensus       127   ~~    128 (129)
T 5HYN_S          127   EM    128 (129)
T ss_dssp               HC
T ss_pred               hc
```

#### HHPRED full length

##### Alignments

Template alignment | NCBI Fasta

1. **XP_006721857.1 PREDICTED: polycomb protein SUZ12 isoform X1 [Homo sapiens]**
Probability: 74.91%, E-value: 21, Score: 47.43, Aligned cols: 71, Identities: 23%, Similarity: 0.453, Template Neff: 5.4

```
Q ss_pred                   cchhhheeeeeccCCCccceeeecccccccCccccccccchhhhhhhHHcc-cceEEEEEeCCceeEEEECCCe
Q Q_8198390          860    PRILSLFYIFKDVNGNVTQKKIVDGWKCGLCGLEEDVGDLNDMSELTVHIKHYH-KYECAVLISGTGVEMVITLPDC   935 (1595)
Q Consensus          860    pril~l~yifkdv~gn~tq~kivdgwkcglcgleedvgdlndmsel~vhikhyh-kyecavlisg~g~emvitlpdc   935 (1595)
                            +..+.++|.|++-+--..|-..++++.|..|++.-     ..|..|..|.+..| .|...+..+-.|+.+.+.+-.|
T Consensus          287    ~~~~~V~y~F~y~nn~~qqte~~dd~~CP~C~~~c-----~sl~~L~~HL~s~H~rF~F~~~~~~~~g~~I~Vsv~e~   358 (603)
T XP_006721857.1     287    RQKLRIFYQFLYNNNTRQQTEARDDLHCPWCTLNC-----RKLYSLLKHLKLCHSRFIFNYVYHPKGARIDVSINEC   358 (603)
T ss_pred                   ccceEEEEEeccCCCccceEEEeCCccCCCccccC-----ccHHHHHHHHHhccccEEEEEEEcCCCCEEEEEecCC
```

Template alignment | Template 3D Structure | PDBe

2. **6C24_A Polycomb protein SUZ12; polycomb repressive complex, AEBP2, JARID2, histone modification, GENE REGULATION; HET: M3L; 3.5A {Hom sapiens}**
Probability: 73.54%, E-value: 23, Score: 48.02, Aligned cols: 68, Identities: 24%, Similarity: 0.493, Template Neff: 5.5

```
Q ss_pred                   hhheeeeeccCCCccceeeecccccccCccccccccchhhhhhhHHcc-cceEEEEEeCCceeEEEECCCe
Q Q_8198390          863    LSLFYIFKDVNGNVTQKKIVDGWKCGLCGLEEDVGDLNDMSELTVHIKHYH-KYECAVLISGTGVEMVITLPDC   935 (1595)
Q Consensus          863    l~l~yifkdv~gn~tq~kivdgwkcglcgleedvgdlndmsel~vhikhyh-kyecavlisg~g~emvitlpdc   935 (1595)
                            +.++|-|+.-++..-|-..+++|.|..|++     +-..+..|..|.+-.| .|........+..|+-+.|++.+|
T Consensus          426    ~~V~Y~F~y~nn~~QqTe~~edf~CP~C~~~~-----~c~sl~~L~~HLk~sH~rF~F~~~~~~~~g~~I~VSin~~   494 (739)
T 6C24_A             426    LRIFYQFLYNNNTRQQTEARDDLHCPWCTL-----NCRKLYSLLKHLKLCHSRFIFNYVYHPKGARIDVSINEC   494 (739)
T ss_dssp                   CCCBCCEESTTTSCCCCCCCSSSSCTTTCC-----CCCSHHHHHHHHHHTTCSEEEECCCTTTTCBCEEEECSC
T ss_pred                   eEEEEEEEeCCceeeEEEEeCCccCCccccC-----cchhHHHHHHHHHHcccCeEEEEEEcCCccEEEEEeccc
```

Template alignment | NCBI Fasta

3. **NP_001308136.1 polycomb protein SUZ12 isoform 2 [Homo sapiens]**
Probability: 73.31%, E-value: 24, Score: 47.13, Aligned cols: 71, Identities: 21%, Similarity: 0.423, Template Neff: 6.7

```
Q ss_pred                   cchhhheeeeeccCCCccceeeecccccccCccccccccchhhhhhhHH-cccceEEEEEeCCceeEEEECCCe
Q Q_8198390          860    PRILSLFYIFKDVNGNVTQKKIVDGWKCGLCGLEEDVGDLNDMSELTVHIKH-YHKYECAVLISGTGVEMVITLPDC   935 (1595)
Q Consensus          860    pril~l~yifkdv~gn~tq~kivdgwkcglcgleedvgdlndmsel~vhikh-yhkyecavlisg~g~emvitlpdc   935 (1595)
                            ...+.++|.|..-+--..|...++++.|..|++.     -..+..|..|.+. +..|.+.+...-.|+.+.+.+-.|
T Consensus          400    ~~~~~v~Y~F~y~~~~~q~te~~~~~~Cp~C~~~~-----c~sl~~L~~HL~~~H~rF~F~~~~~~~~g~~I~Vs~~~~   471 (716)
T NP_001308136.1     400    RQKLRIFYQFLYNNNTRQQTEARDDLHCPWCTLN-----CRKLYSLLKHLKLCHSRFIFNYVYHPKGARIDVSINEC   471 (716)
T ss_pred                   ccceEEEEEecCCCCcceEEEeCCccCCccccCC-----ChhHHHHHHHHHhccccEEEEEEEcCCCCEEEEEeccC
```

Fig.S3 Cryptococcus Bnd1 = SUZ12. Shown are the alphafold determined structures of Cryptococcus Bnd1 and Homo sapiens SUZ12. In addition we here show the HHPRED outputs of a search with the VEFS box of Cryptococcus and an HHPRED output of a full length search with Cryptococcus SUZ12

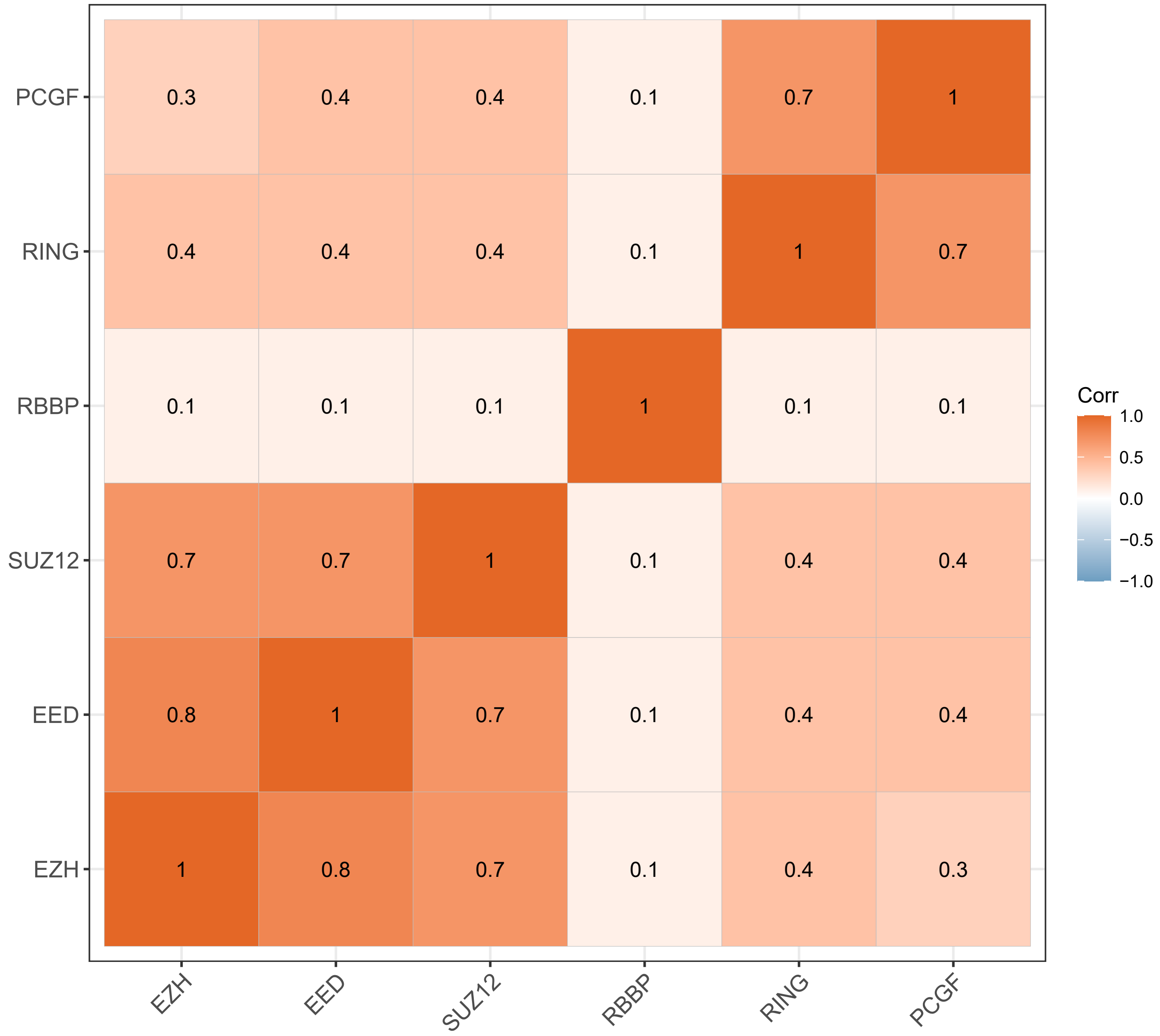


Fig.S4. Heatmap of pearson correlation scores. Intra-subunit correlation is higher than inter-subunit correlations

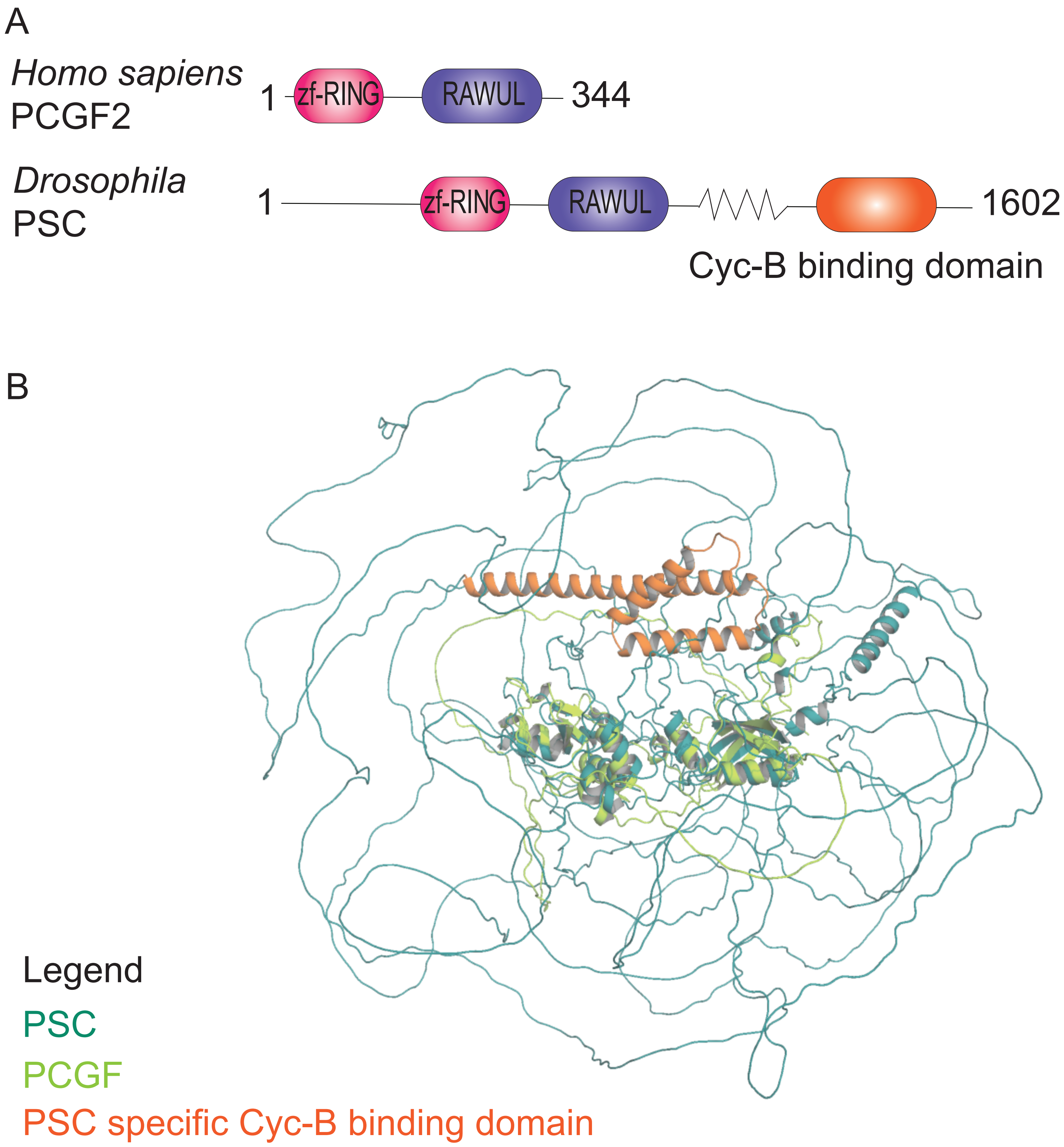


Fig.S5 Diptera-specific domain to bind and regulate cyclin B to control cell cycle progression. (A) Schematic overview of Homo sapiens PCGF2 and Drosophila PSC with the Cyc-B binding domain highlighted in orange. (B) Alignment of Alphafold structures of Homo sapiens PCGF2 (light green) and Drosophila PSC (teal). Highlighted in orange is the Cyc-B binding domain.
